# Generic assembly patterns in complex ecological communities

**DOI:** 10.1101/145862

**Authors:** Matthieu Barbier, Jean-Franois Arnoldi, Guy Bunin, Michel Loreau

## Abstract

The study of ecological communities often involves detailed simulations of complex networks. But our empirical knowledge of these networks is typically incomplete, and the space of simulation models and parameters is vast, leaving room for uncertainty in theoretical predictions. Here, we show that a large fraction of this space of possibilities exhibits generic behaviors that are robust to modelling choices. We consider a wide array of model features, including interaction types and community structures, known to generate different dynamics for a few species. We combine these features in large simulated communities, and show that equilibrium diversity, functioning and stability can be predicted analytically using a random model parameterized by a few statistical properties of the community. We give an ecological interpretation of this “disordered” limit where structure fails to emerge from complexity. We also demonstrate that some well-studied interaction patterns remain relevant in large ecosystems, but their impact can be encapsulated in a minimal number of additional parameters. Our approach provides a powerful framework for predicting the outcomes of ecosystem assembly and quantifying the added value of more detailed models and measurements.

## Introduction

Ecological communities form large and intricate networks of dynamical interdependencies between their constituent species and abiotic factors. The richness and variety of natural systems has long been a source of wonder, but understanding the mechanisms that shape their diversity, stability and functioning is also a pressing challenge [26, 19, 6, 25]. Their theoretical investigation often relies on extensive numerical simulations modelled after, but not fully parameterized by, empirical observations, e.g. [10]. Although current computational power allows exploring levels of complexity that were unapproachable only a few decades ago, the parameter space in these simulations is fatally large, with numerous plausible choices for species traits, interactions, and even dynamical equations, that may give different and even contradictory predictions.

Against this vertiginous perspective, we reveal emergent *genericity* allowing complex interaction networks, despite their superficial differences, to be understood within a common framework and distinguished by only a few parameters. This is possible because complex structure does not always entail complex dynamics. In the setting of ecological assembly, we show that various community models display similar collective behaviors, which can be captured using four universal parameters. These parameters have recently been identified in a random Lotka-Volterra system [11] interpreted here as the “disordered limit” for a broad class of models. We use this random limit to make null predictions for community properties, which we compare to the outcomes of specific models to quantify the added value of their structure. When models depart from null expectations, we propose ways to expand the random model to capture relevant features while retaining maximal genericity.

This corroborates a long-standing intuition in ecology: simple laws can emerge from the interaction of many species. Indeed, ecologists have long posited that networks can be reduced to selected features [31, 3], or to averaged-out [20] or random interactions [29, 40]. On the other hand, recent approaches, starting from complex models, have attempted to demonstrate this emergent simplicity [16] but their success has been restricted to specific interaction types [1]. We thus bridge the gap between these seminal works by developing a reductive approach that is applicable to a wide array of complex systems, and that can be used to identify relevant structures, allowing to evaluate previous simplified theories.

To clearly see the emergence of genericity in ecological interactions, we must first set aside other important drivers of community dynamics, such as stochasticity and external perturbations. Instead, we focus here on the *stable* and *uninvadable* equilibria that result from community assembly, as they are determined by species traits and interactions only. We consider models of community assembly within the class of generalized Lotka-Volterra dynamics

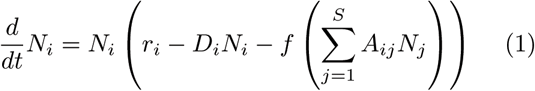

where *i* runs over a pool of *S* species, with *N*_*i*_, *r*_*i*_ and *D*_*i*_ their abundance, intrinsic growth rate and self-regulation (density-dependent mortality) respectively, *ƒ* the functional response [18], and *A*_*ij*_ the per-capita interaction coefficients. Models differ by the structure of their coefficients and by their assembly process - how and when species from the fixed regional pool may invade the community [24].

We propose that generic behaviors can be identified by comparing these models to their *disordered limit*, obtained by randomizing the effective “carrying capacities” and interactions

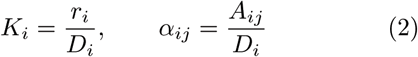

while preserving some essential statistics:

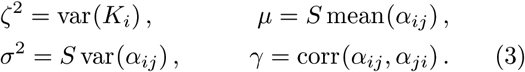

It was shown in [11] that these four quantities - hereafter carrying capacity spread *ζ*, interaction antagonism *μ*, heterogeneity *σ* and reciprocity *γ* - emerge as the only relevant parameters in large random systems^1^. From these parameters, analytical formulas provide the typical abundance, diversity and stability properties of uninvadable equilibria, regardless of the assembly process [11, 9].

As illustrated in Fig. 1, we draw from the literature [27, 13, 17, 5, 23] a list of important model features, including interaction types (mutualism, competition and predation), community structures, functional responses, and parameter distributions (random or mechanistically derived). We then combine these features and vary their parameters to generate many distinct communities. We simulate their dynamics from random initial conditions until an unin-vadable community is reached, and measure its stability, diversity and functioning (see (6) in Materials and Methods). We finally compare these models to their disordered limit as defined above, hereafter the *reference model*, to obtain null predictions for the measured properties.

**Figure 1:**
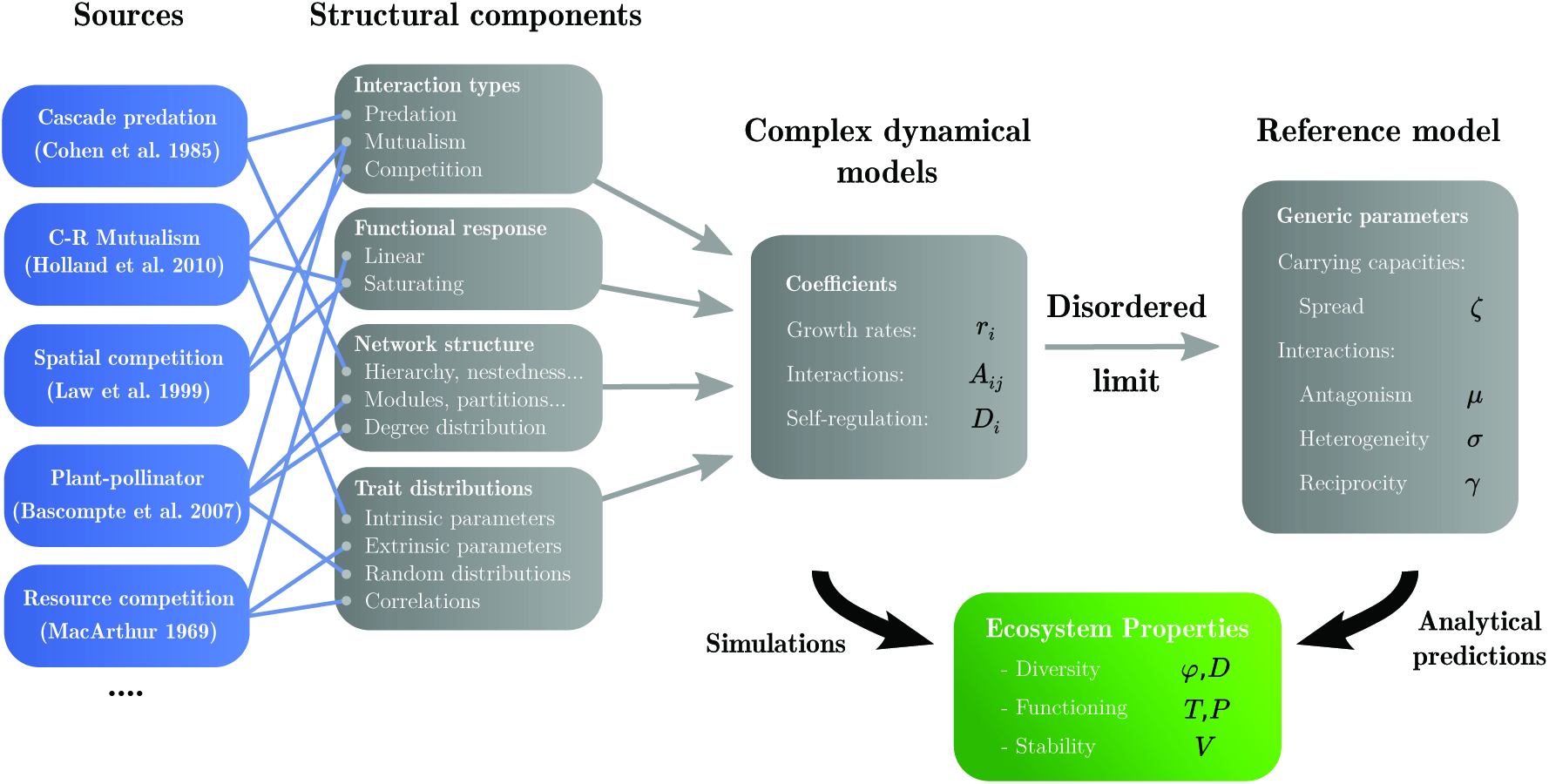
General outline of our approach. On the left, examples of sources from the literature, from which model features have been extracted (see full list in SI Appendix, Sec. II.1). Diverse combinations of these model features and variations of their parameters yield distinct communities, characterized by the coefficients and functional response in the dynamics (1). We simulate them until they reach an assembled equilibrium, whose properties we measure. We then randomize interactions and carrying capacities, preserving the four statistics in (3). The randomized community’s equilibrium properties are known analytically from the solution of the reference model [11] and can be compared to simulation outcomes, see Fig. 2.

Where predictions agree with simulation results within 5% according to the error metric in (7), we judge that a model’s behavior is well-predicted by its disordered limit, meaning that the generic parameters in (3) capture all dynamically-relevant features. In case of a discrepancy, we demonstrate that it is possible to recover quantitative agreement through a minimal addition of structure: either extracting more precise statistical information such as correlations (see SI Appendix IV), or dividing the community into a few modules and computing parameters within and between these groups. In both cases, we are still able to reduce a complex model to a limited number of parameters, which does not grow with network size or connectance.

## Results

In Fig. 2, we showcase some of the predicted community properties in a particular example: a resource competition model (see Materials and Methods). We compare simulation results to analytical predictions in the disordered limit, i.e. the reference random model parameterized by a few aggregate statistics, see (3). Despite lacking any underlying mechanistic structure, this random model reproduces all patterns *quantitatively*, demonstrating that the structure and coefficients of resource competition are significant only inasmuch as they affect the reference parameters.

**Figure 2:**
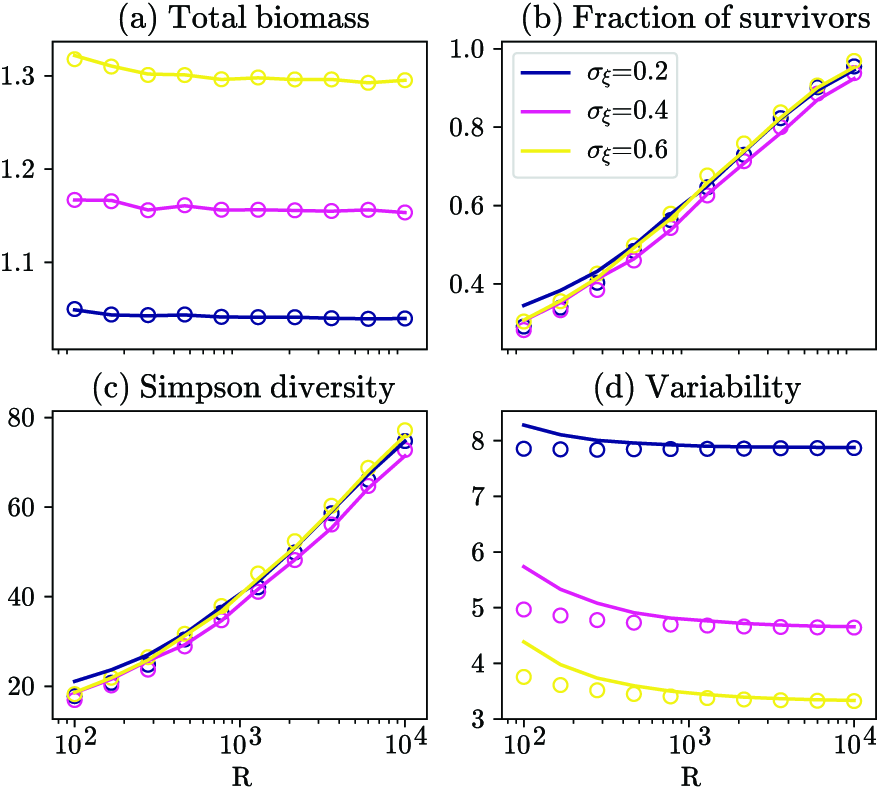
Simulation results for the resource competition model (dots) and analytical predictions in the disordered limit (lines) for various community properties: (a) Total biomass *T*, (b) Fraction of surviving species *ϕ*, (c) Simpson diversity *D*, (d) Temporal variability *V*. We vary the number of resources *R* and the heterogeneity of consumption rates *σ*_*ξ*_ (see Materials and Methods). This then affects the value of the generic parameters in (3), inserted into the analytical solution of the reference model to obtain null predictions. The remarkable agreement confirms that the resource competition model exhibits fully disordered behavior.

We illustrate in Fig. 3 how different models visit this common parameter space as we vary their specific properties. For instance, models can overlap: we construct an example where competition and predator-prey models correspond to the same values of all four parameters, giving rise to communities with identical properties – in particular, species abundance distributions. These very different models thus live in the same space of generic dynamics. However, they mostly occupy different regions of this space, and overlap is rare for all four parameters: for instance, resource competition tends to lead to higher antagonism *μ* but lower heterogeneity *σ* than preda-tion, entailing lower total biomass but more stability.

**Figure 3:**
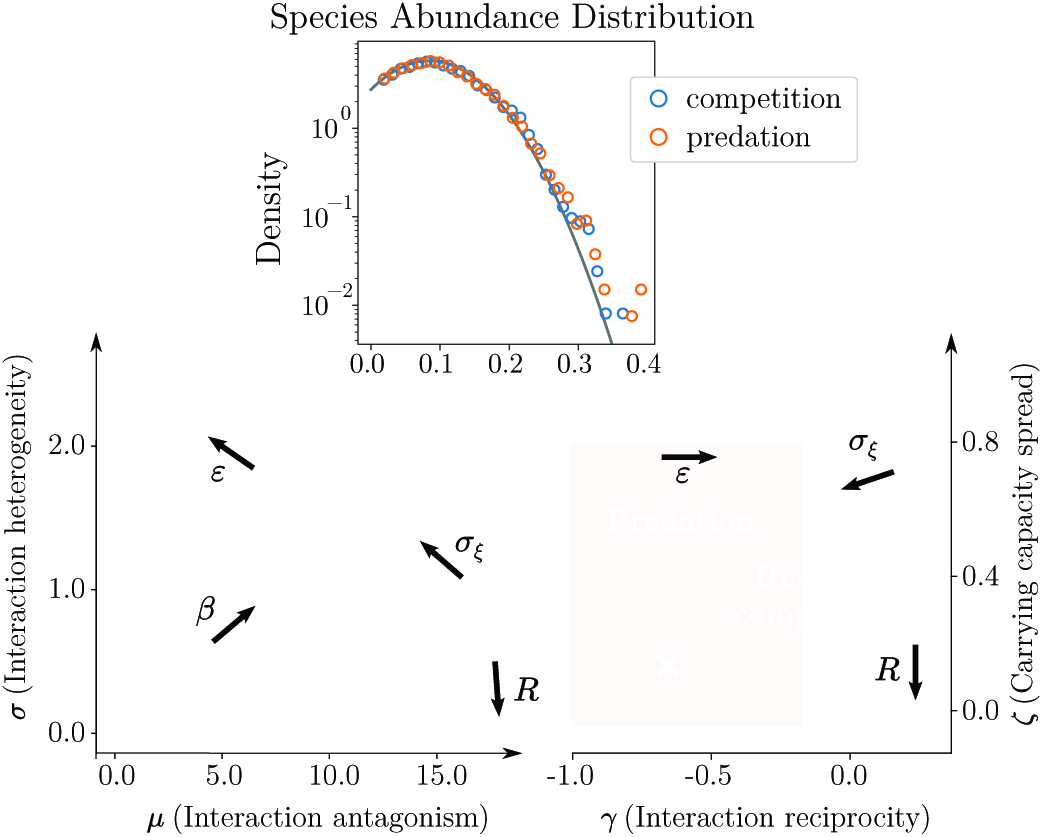
Different models visit the generic parameter space (*μ, σ, γ, ς*) defined in (3) as we vary their model-specific control parameters (bold arrows, defined in Materials and Methods). We illustrate in the (*μ, σ*)> and (*γ, ς*) planes the regions visited by two models. In orange, predation with mean intensity *β* ∈ [0.1,25] and conversion efficiency *ε* ∈ [0,1] (in this model, *ς* is a free parameter). In blue, resource competition with resources number *R* ∈ [10^2^,10^4^] and consumption heterogeneity *σ*_*ξ*_ ∈ [0.1,0.6]. *Inset:* An example where a competitive community and a predator-prey community display identical species abundance distributions, well-predicted by the theory (solid line), corresponding to the parameter values marked by the cross. We discuss below and in SI Appendix (Sec. III.3) under which conditions these distributions are predicted to be narrow (e.g. normal) or fat-tailed (e.g. lognormal), see also [40, 41].

While the reference model has significant unifying and explanatory power, it only represents a limit in which all system-level structure is erased. We next investigated which model assumptions among those in Fig. 1 caused a deviation from this disordered limit. We show in Fig. 4 that many choices failed to yield any deviation^2^. We plotted the relative error of the analytical predictions compared with simulations (see (7), Materials and Methods) as we added more and more structure to the interactions. Strikingly, pure mutualistic communities were found to be uniquely disordered^3^: while their network structure has been extensively studied [4, 35], we found that it did not contribute to global community properties, except indirectly by changing the reference parameters in (3). By contrast, we found that some structural properties – cascade, nestedness and partitions – led to different dynamical behavior in other interaction types. Structure strongly mattered in competitive communities, even though they are the least studied in that respect, except for hierarchies [33]. In particular, a notable deviation from disorder occurred in the bipartite case, i.e. competition between two groups of freely coexisting species (e.g. an idealization of a contact zone between communities). Even with weak pairwise interactions, we observed that one entire group could go extinct while the other survived, suggesting group-level competitive exclusion [39].

**Figure 4:**
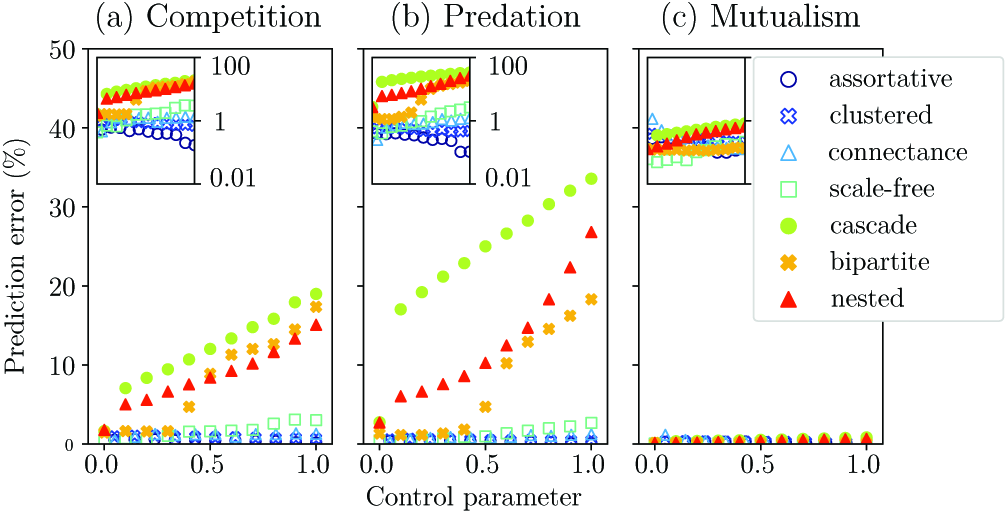
Network structure and deviation from the disordered limit. (a)-(c) For the three main interaction types, we show the relative error (y-axis, between 0 and 50%) of the reference model predictions against simulations, see (7). *Inset*: Same results in log scale. The symbol sets correspond to different network structural properties: assortativity, partitioning, clustering, nestedness and scale-free or cascade structure. Each comes with a specific control parameter (x-axis, see list in SI Appendix, Sec. II.1.2), allowing to go from an Erdos-Renyi random graph to a maximally structured network. For instance, the probability *p*_*d*_ of attaching preferentially to nodes with higher degree yields a scale-free network when *p*_*d*_ = 1. We also vary connectance in a random graph. We see that only bipartition, cascade structure and nestedness cause deviations from null predictions in competitive and predatory interactions, and none do in mutualistic communities (where interaction strength is limited, see Materials and Methods).

Thus, various examples including nestedness, partitions and functional groups demonstrate how complex models deviate from fully disordered dynamics. Additional information is then needed to predict community properties. We illustrate in Fig. 5 the case of a plant-pollinator community with inter-group mutualism and intra-group competition. A bipartite mutualistic community exhibits fully disordered behavior (Fig. 4) but adding intra-group competition caused results to deviate from the disordered limit. By extending our theory to distinguish multiple groups and computing interaction parameters within and between groups, we made accurate predictions, even in the complex situation of intermediate levels of order, where the boundaries between groups were blurred. Similarly, we describe in SI Appendix (Fig. S4) how other structures such as hierarchies can be accounted for, and note (Fig. S3) that a saturating functional response makes these structures less relevant. These results suggests the value of incorporating simple structure and disorder simultaneously in a model to tackle more complex communities.

**Figure 5:**
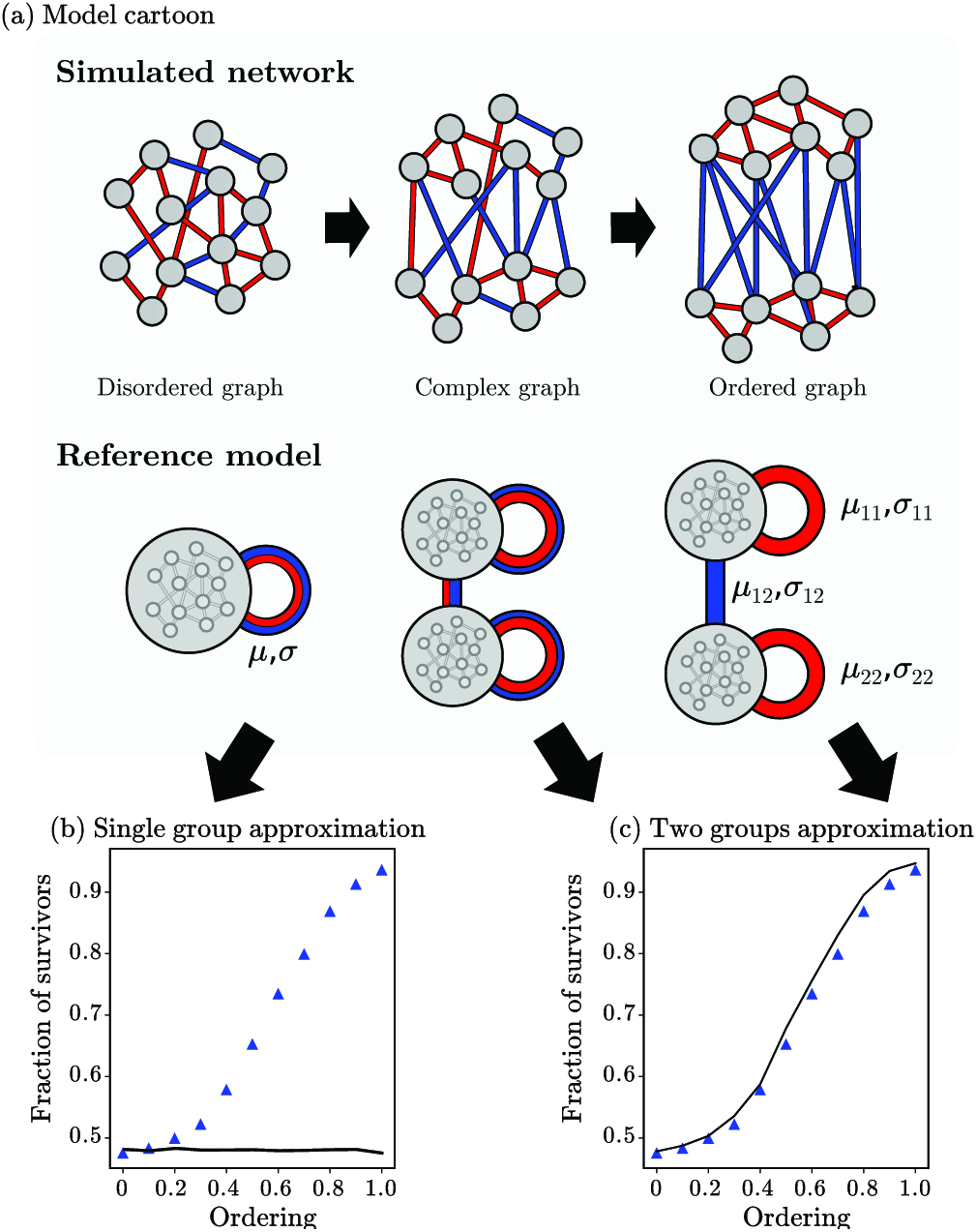
Genericity beyond full disorder. We give an example of structure: two functional groups with competitive and mutualistic interactions. (a) Cartoon of the model. In the ordered limit, all intra-group interactions are competitive and all inter-group interactions are mutualistic. The ordering parameter is the probability to rewire interactions without respecting group structure. (b) Fraction of surviving species in the assembled community for the simulation model (symbols) against analytical predictions in the disordered limit (solid line) which cannot account for ordering. (c) Same data, but the reference model is extended with more structure, distinguishing between parameters for inter-and intra-group interactions. It successfully predicts community properties for any degree of ordering, even the complex intermediate case where group boundaries are blurred.

## Discussion

By comparing dynamical ecological models to their disordered limit, i.e. a randomization preserving only a few statistics of species traits and interactions, we provide a novel approach to study the emergence of simple and generic patterns in community assembly. Dynamics in the disordered limit are described in a common framework, a random Lotka-Volterra *reference model*, within which community properties (diversity, stability and functioning) can be predicted analytically [11] provided that interactions are distributed among many species. As these predictions match simulation outcomes for a wide array of complex models, we conclude that, although superficially different, these models exhibit the same generic behavior. Some large-scale community structures, however, including partitions, nestedness, hierarchy (Fig. 4) and functional groups (Fig. 5), do induce a discrepancy between simulation models and their disordered limit. Taking this structure into account via a few additional parameters, we obtain good agreement from a less generic, but still greatly simplified analytical model.

### Disorder and genericity

The significance of our findings hinges on two questions: why this choice of disordered limit and generic parameters, and what is their ecological meaning? First, let us highlight a feature of the random reference model: its many-species deterministic dynamics can be mapped to, and solved as, a single stochastic equation [32, 11]. In other words, it only captures the most generic collective dynamics (see SI Appendix, Fig S1) because it effectively describes a sin gle stochastic pseudo-species, with an internal distribution of abundances and interactions. This can easily encompass other ecological heterogeneities, such as variation between individuals or populations.

Our work thus shows that, unlike a model with a single deterministic species [16] or many identical ones [20], a model with a single stochastic pseudo-species may in fact have the same richness of behavior as a complex community. Our extension to functional groups suggests that a more complex community can be represented by a few interacting stochastic pseudo-species, without loss of information. This dynamical richness is captured by the generic parameters in (3), especially *σ* and *γ* whose role is more subtle: as *σ* increases, species interactions become less similar, hence more likely to create shifting conditions favoring some species, reducing both diversity and stability. In turn, *γ* represents interaction symmetry, negative for trophic systems as well as asymmetrical competition or mutualism. This causes a negative feedback of a species on itself via its partners, leading to stabler, more species-rich equilibria [11].

We find a general criterion for this disordered limit: heterogeneities in species traits and interactions should be well-mixed throughout the community, so that each species and its neighborhood constitute an unbiased (even if partial) sample of the whole system. This criterion holds true regardless of many model details, including interaction type, functional response and diverse network metrics (connectance, assortativity, clustering). As expected from this intuitive criterion, we found that structures that lead to different outcomes than the reference model must exhibit large-scale differentiated neighborhoods in the web of interactions, e.g. functional groups or different positions within a hierarchy.

### Theoretical consequences

Our results have at least two consequences for the theory of complex ecological communities. On the one hand, theoretical investigations aiming for generic patterns can start from the reference model as a flexible and simple platform for exploring various ecological patterns. If a system satisfies our conditions for generic behavior, it is enough to know how its parameters and assumptions translate into the reference parameters in (3) to deduce their consequences on the assembled state.

On the other hand, theorists interested in a more specific and complex model may benefit from comparing its predictions to that of the reference model. This is immediate as there is no fitting involved: the parameters of the reference model are simple statistics of the species pool, and can be inserted into our analytical solution to readily obtain predictions. Checking when these results align is a way of testing the “added value” of other structures and mechanisms. The extra information needed to reconcile simulations with the reference model, if any, provides a measure of a community’s effective complexity: for instance, how many functional groups must be inferred and parameterized for an extended theory (as in Fig. 5) to make accurate predictions on a complex network.

The picture that emerges is that complexity peaks at intermediate levels of heterogeneity (see SI Appendix, Fig. S1), a relationship found across domains [36, 7]. A bipartite network distinguishes clear neighborhoods, but a highly multipartite one, such as an intricate web of functional groups and fluxes, may again tend to resemble a disordered set of species. Many additional details, including network metrics, degree distributions or mechanistic parameterization, contribute to species heterogeneity but fail to add true complexity.

### Empirical consequences

From an empirical perspective, the details of species interactions are often difficult to observe and measure [8]. That relevant ecological patterns may only depend on aggregated properties is a positive message: we could make predictions that are robust to a lack of detailed information, and that rely on a minimal number of fitted or inferred parameters [14]. We expect this approach to succeed when each species interacts with a fair sample of the overall community, e.g. when interactions are “well-mixed” due to spatial heterogeneity (competitive plant communities) or mediated by a common resource (microbial public goods).

More generally, multitrophic ecological communities such as plant-pollinator or plant-herbivore communities could be understood by a combination of simple structure and disorder. Much empirical work has focused on evidencing salient interaction patterns through the examination of particular model species and their pairwise functional relationships. We propose that these insights could be combined with more aggregated data to construct models, such as our bipartite example (Fig. 5), that encapsulate both the essential functional structure and the diversity and complexity of a realistic community, with few parameters to estimate. Coherent global structure has indeed been found in large-scale empirical studies involving thousands of interactions, showing a limited set of distinct interaction patterns [21]. Our suggested approach may be seen as a step towards combining local and global, species and ecosystem perspectives.

### Implications for future work

We have shown that, even when we try to build a detailed picture of an ecological community, its collective dynamics can often be understood from a few large-scale properties, that do not always follow the intuitive categories of ecological mechanisms. Our work offers an outlook on what complexity means in an ecological setting. Predicting the fate of a certain species at a given locale, e.g. for conservation, may require knowledge about every important feedback within its environment, biotic and abiotic. But one rarely needs all these details at once to understand the aggregate properties of an ecosystem, or the fate of most species most of the time. Instead, the exhaustive study of ecological networks could also pave the way toward finding new dimensions of simplicity at the collective level.

The idea of “disorder” does not restrict our approach to models with random interactions. Purely random systems are in some sense special – the total absence of order is a peculiar feature in the infinite space of possible communities. Instead, our results suggest that predictions from a random model succeed wherever non-random motifs interfere due to their number, heterogeneity and mixing. Combining disorder and coherent structure to understand complex objects has deep mathematical underpinnings: Terence Tao speaks of “ a fundamental dichotomy between structure and randomness, which in turn leads (roughly speaking) to a decomposition of any object into a structured (low-complexity) component and a random (discorrelated) component.” [38].

While we have focused on equilibrium properties, the methods we employed can extend to full dynamics, including stochasticity [32], higher-order interactions [15], and even evolutionary dynamics [34]. In particular, a connection to other theories of assembly, e.g. [28], could be made by considering open ecosystems or metacommunities with explicit immigration processes, which could also lead to more realistic abundance distributions [41].

Finally, we must address the general biological relevance of these findings. Many ecologists have taken sides in a famous theoretical and empirical debate: whether biodiversity allows for stable ecosystems [29, 30]. Our work suggest a broader claim, encompassing both sides: biodiversity allows for stable *laws*, i.e. predictable relationships between various ecosystem properties such as stability, functioning and diversity. These relationships may be idiosyncratic in small systems, but the more species-rich and heterogeneous a community, the more it tends toward a generic limit (see SI Appendix, Fig S1) where many of its properties covary in a universal parameter space.

Could these generic patterns be precluded or favored by evolution, which may either foster specialized structures, or exploit the universality and robustness of disorder? Understanding these interactions with adaptive and evolutionary processes should be at the heart of future developments.

## Methods

### Simulation models

All simulation models are detailed in SI Appendix II, with example models from the figures in main text described in Table S1 and Sec. II.1.5. For resource competition (Fig. 2 and 3), dynamical coefficients are computed from the abundance *ρ*_*a*_ of resource *a* and its consumption rate *ξ*_*ia*_ by species *i*, both drawn as i.i.d. random variables with variances 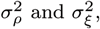, as

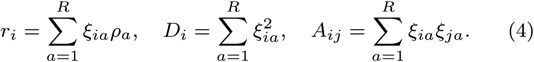

For the predation model (Fig. 3) we randomly select within each pair of species (*i, j*) a prey *i* and predator *j*, draw *β*_*ij*_ as i.i.d. exponential variables with mean *β*, and define

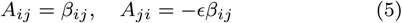

with ε the biomass conversion efficiency. No trophic ordering is imposed in the species pool. Finally, for mutualistic interactions with a linear functional response (Fig. 4), interactions were rescaled so that Σ_*j*_|*A*_*ij*_| ≤ 0.5*D*_*i*_ to prevent population explosion.

### Community properties

An R implementation of the algorithm used to compare simulation results to predictions is maintained at http://github.com/mrcbarbier/ecocavity-R, with example files.

Properties of the assembled community were measured in simulations, and derived analytically in the reference model (see details in SI Appendix, Sec. III.4). We focused here on five properties:

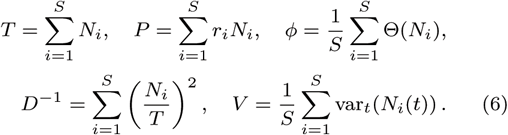

with *Θ*(*N*_*i*_) = 1 if *N*_*i*_ > 0, 0 otherwise. Ecosystem functioning is characterized by total biomass *T* and total productivity from external resources *P*. Diversity is represented by two quantities: *ϕ*, the fraction of species that survive in the assembled state, and *D*, the inverse of Simpson’s index [37]. Among the many dimensions of ecological stability, we focus on empirically-relevant variability *V*, the variance in time of species abundance due to stochastic perturbations [22, 2]. Finally, all these quantities were combined into a single metric of relative error of the analytical predictions:

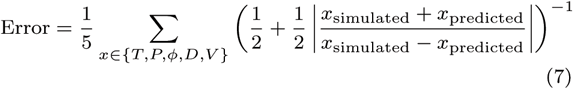

which is symmetrical and has values in [0, 1].

### Master equation for the stochastic pseudo-species

In the disordered limit, (1) can be transformed into a single implicit integral equation (see SI Appendix III) for the abundance distribution *P*(*N*), given the distribution *p*(*K*) of carrying capacities:

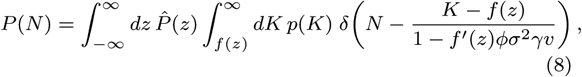

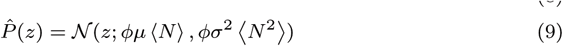

where *𝒩*(*x;m, s*^2^) denotes the normal distribution of *x* with mean *m* and variance *s*^2^, and the state variables *ϕ*, 〈*N*〉, 〈*N*^2^〉 and *v* are computed implicitly through the moment equations

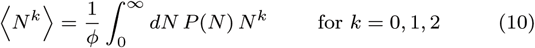

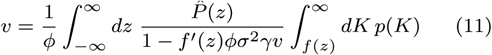

which can be integrated numerically (see SI Appendix, Numerical solution of the reference model). If *ƒ* is linear and *p*(*K*) = *𝒩*(*K*; 1, *ζ*^2^), then *P*(*N*) is a truncated Gaussian [11] as seen in Fig. 3.

## Acknowledgments

We address our thanks to Bart Haegeman for insightful discussions during the preparation of this work, and Sbastien Ibanez, Sonia Kfi and four anonymous reviewers for many helpful comments on the manuscript. This work was supported by the TULIP Laboratory of Excellence (ANR-10-LABX-41) and by the BIOSTASES Advanced Grant, funded by the European Research Council under the European Unions Horizon 2020 research and innovation programme (666971).

## Footnotes

1 More precisely, the effects of species interactions become truly generic, and entirely characterized by *μ, σ* and *γ*, in large communities devoid of very strong pairwise interactions (e.g. competitive exclusion). This emergent genericity of interactions does not extend to intrinsic species properties (carrying capacities), which can therefore retain model-specific effects. Their most generic effects are captured by *ς* (see SI Appendix III.2), but some results below require one more parameter, *C*_*Kα*_, the correlation between carrying capacities and interactions, e.g. in a competition-colonization tradeoff [12] or in resource competition (see SI Appendix II.1.5). The random model remains fully solvable for any distribution of intrinsic species traits.

2 Here and in Table S2 in SI Appendix, we systematically explore combinations of one interaction type and another model feature – trait distributions or network structures. All results in Fig. 4 were reproduced with a saturating functional response, see Fig S3. More complex combinations were not tried systematically, but some figure as individual examples, see SI Appendix Table S1.

3 First, mutualistic interactions must either be weak, or have a saturation threshold, to prevent population explosion, which drastically limits their heterogeneity. Second, in the absence of any negative interaction, the community can effectively be reduced to a single variable [16, 1].

